# PhosIDP: a web tool to visualize the location of phosphorylation sites in disordered regions

**DOI:** 10.1101/2021.04.13.439614

**Authors:** Sonia T. Nicolaou, Max Hebditch, Owen J. Jonathan, Chandra S. Verma, Jim Warwicker

**Affiliations:** School of Biological Sciences, Faculty of Biology, Medicine and Health, Manchester Institute of Biotechnology, University of Manchester, Manchester M1 7DN, UK; Bioinformatics Institute, Agency for Science, Technology, and Research (A*STAR), Singapore 138671, Singapore; School of Biological Sciences, Nanyang Technological University, 60 Nanyang Drive, Singapore 637551; Department of Biological Sciences, National University of Singapore, 14 Science Drive 4, Singapore 117543

## Abstract

Charge is a key determinant of intrinsically disordered protein (IDP) and intrinsically disordered region (IDR) properties. IDPs and IDRs are enriched in sites of phosphorylation, which alters charge. Visualizing the degree to which phosphorylation modulates the charge profile of a sequence would assist in the functional interpretation of IDPs and IDRs. PhosIDP is a web tool that shows variation of charge and fold propensity upon phosphorylation. In combination with the displayed location of protein domains, the information provided by the web tool can lead to functional inferences for the consequences of phosphorylation. IDRs are components of many proteins that form biological condensates. It is shown that IDR charge, and its modulation by phosphorylation, is more tightly controlled for proteins that are essential for condensate formation than for those present in condensates but inessential.

## Introduction

Intrinsically disordered proteins (IDPs) and intrinsically disordered regions (IDRs) of proteins have been challenging the traditional structure-function paradigm over the last two decades. They exhibit a range of conformations, from molten globules to random coils ^1^, with their net charge correlated to their conformational preference ^2,3^. The absence of distinct structure in IDPs/IDRs can be attributed to their high net charge and low hydrophobicity ^4^. The flexible nature of IDPs/IDRs allows them to be easily regulated by post-translational modifications (PTMs) ^5^. Phosphorylation is a post-translational modification enriched in IDPs/IDRs ^6^, that alters the charge of serine, threonine and tyrosine amino acids by substituting a hydroxy functional group with a negatively charged phosphate group. Phosphorylation plays a crucial role in many biological processes, where it can modify charge and hydrophobicity, and modulate interactions with partners ^7^.

According to the polyelectrostatic model, interconverting conformations of charged IDPs and/or IDRs bind to their partners through an average electrostatic field caused by long-range electrostatic interactions ^8–10^. Phosphorylation can modify the electrostatic interactions of disordered regions by altering their charge which can either enhance or reduce their binding affinity for binding partners ^10^. Conformational ensembles of IDPs/IDRs depend on the balance of all charge interactions in the protein, rather than individual short-range interactions between the binding site of the protein and a binding partner. Therefore, they can interact with partners through several distinct binding motifs and/or conformations, and various functional elements will switch between availability for interaction and burial ^10^.

It is becoming clear that phase-separating proteins and other molecules, such as mRNA, mediate a variety of biological functions ^11^. Proteins containing IDRs are commonly found in membraneless organelles, formed by liquid-liquid phase separation (LLPS) ^12^. Messenger RNAs are also commonly sequestered during LLPS, typically alongside proteins with IDRs and specific RNA-binding motifs ^12^, so specific and non-specific charge-charge interactions are likely to be involved. Phosphorylation (and therefore charge variation) is a known control mechanism in LLPS formation ^13^. Databases of proteins observed to associate with membraneless organelles are being collated. Notably, the DrLLPS resource ^14^ classifies proteins undergoing LLPS as core (localised proteins that have been verified as essential for granule assembly/maintenance), client (localised but non-essential), and regulator (contribute to modulation of LLPS but are not located in the condensate).

PhosIDP is an addition to the protein-sol software suite ^15^, created for visualizing the effects of phosphorylation on protein sequence charge profiles, and thereby facilitating functional hypotheses. In order to assist in understanding the behaviour of IDPs/IDRs, the web tool visualizes changes resulting from phosphorylation across a sequence, rather than focusing on the consequences of individual phosphorylation sites. Further, offline calculations for datasets of proteins are presented that demonstrate the role of phosphorylation in modulating charge for IDRs in core LLPS proteins, but not client proteins.

## Results

### Phosphorylation mediates substantial charge alteration in the IDR of CIRBP

As an exemplar of proteins of interest, human cold-inducible RNA binding protein (CIRBP, UniProt ^16^ ID Q14011) is used, in particular as a representative of a core set of proteins that are crucial for the formation of stress granules (SGs), examples of liquid-liquid phase separated membraneless organelles ^17^. CIRBP possesses a structured N-terminal RNA recognition motif (RRM) that mediates specific RNA interactions, and a C-terminal disordered arginine/glycine-rich (RGG) region that is involved in weak multivalent RNA interactions ^17^. Both the positive charge and disorder of the unphosphorylated RGG region are apparent in Fig. 1 (panels A and B). It is known that phosphorylation can regulate the phase transitions of stress granules ^13^. Here, the incorporation of phosphorylation (as recorded in UniProt) shows that the C-terminal end of the disordered region of CIRBP becomes substantially more negatively charged, noting that the N-terminal end of the disordered region remains positively charged.

**Figure 1.**
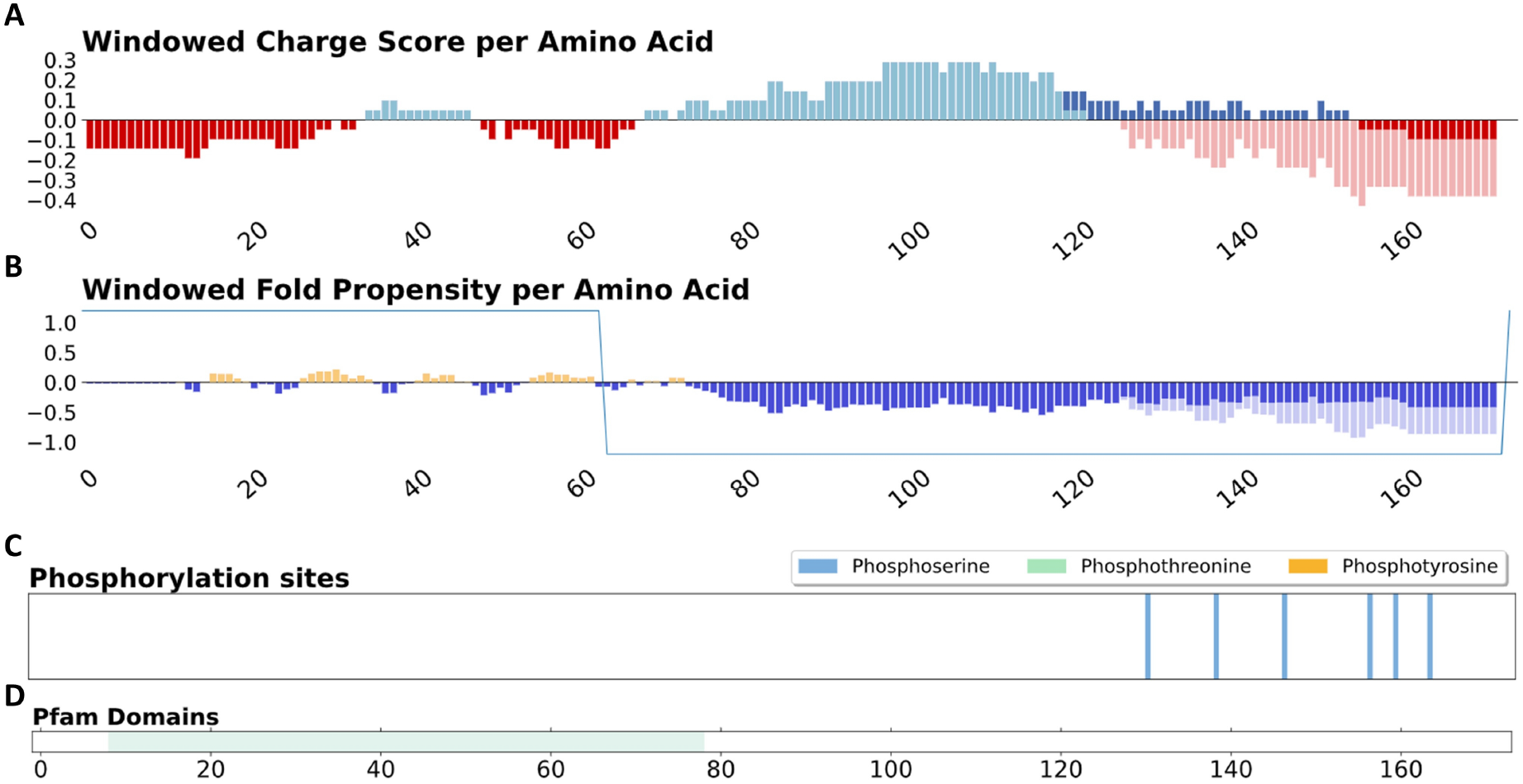
Results of phosIDP server for CIRBP (UniprotKB: Q14011), a stress granule-associated protein. (A) and (B) Changes to charge and fold propensity upon phosphorylation are depicted in lighter shades. The sawtooth window for smoothing disordered and ordered regions is shown in blue. (C) Phosphorylation sites from UniProt are displayed. (D) Location of known Pfam domains.

### Nucleophosmin, a protein with order-disorder transitions coupled to phosphorylation

The potential of our server to highlight the role phosphorylation could play in structural transitions is demonstrated with human nucleophosmin, which (in common with CIRBP) is classified as a core protein in regard to formation of membraneless organelles ^14^. Nucleophosmin shuttles between the nucleus and cytoplasm, undertaking multiple roles ^18^. The current focus is on how prediction of structured and unstructured regions in the phosIDP server, and the presence of phosphorylation sites, correlates with segments for which structural flexibility is known to correlate with function. The core domain is structurally polymorphic, with pentamer formation dependent on phosphorylation status ^19^. Phosphorylation biases towards monomer over pentamer, and this balance is also influenced by ligand binding, contributing to the rich functional properties of nucleophosmin ^20^. Only a part of the folded core domain is predicted as structured, using sequence-based fold propensity (Fig. 2). Colour-coding is common between the predicted structured/unstructured plot (Fig. 2B) and the core domain structure (Fig. 2E, 4n8m ^19^), with the predicted structured and unstructured regions forming the two parts of the monomer. Extensive interactions of the monomer within the pentamer (Fig. 2E) are consistent with the 3D structural stability being dependent upon oligomerisation. Furthermore, the localisation of phosphorylation sites at the monomer interface likely reflects their role in the monomer – pentamer equilibrium ^19^. Therefore, if a user were studying the phosIDP server results, in the absence of known 3D structure, a reasonable prediction would be that conformation predicts only as weakly folded, and that phosphorylation alters the structured/unstructured balance, thereby potentially mediating function. Such a hypothesis could then be subject to the types of experimental analysis that have been applied to investigate the nucleophosmin core domain.

**Figure 2.**
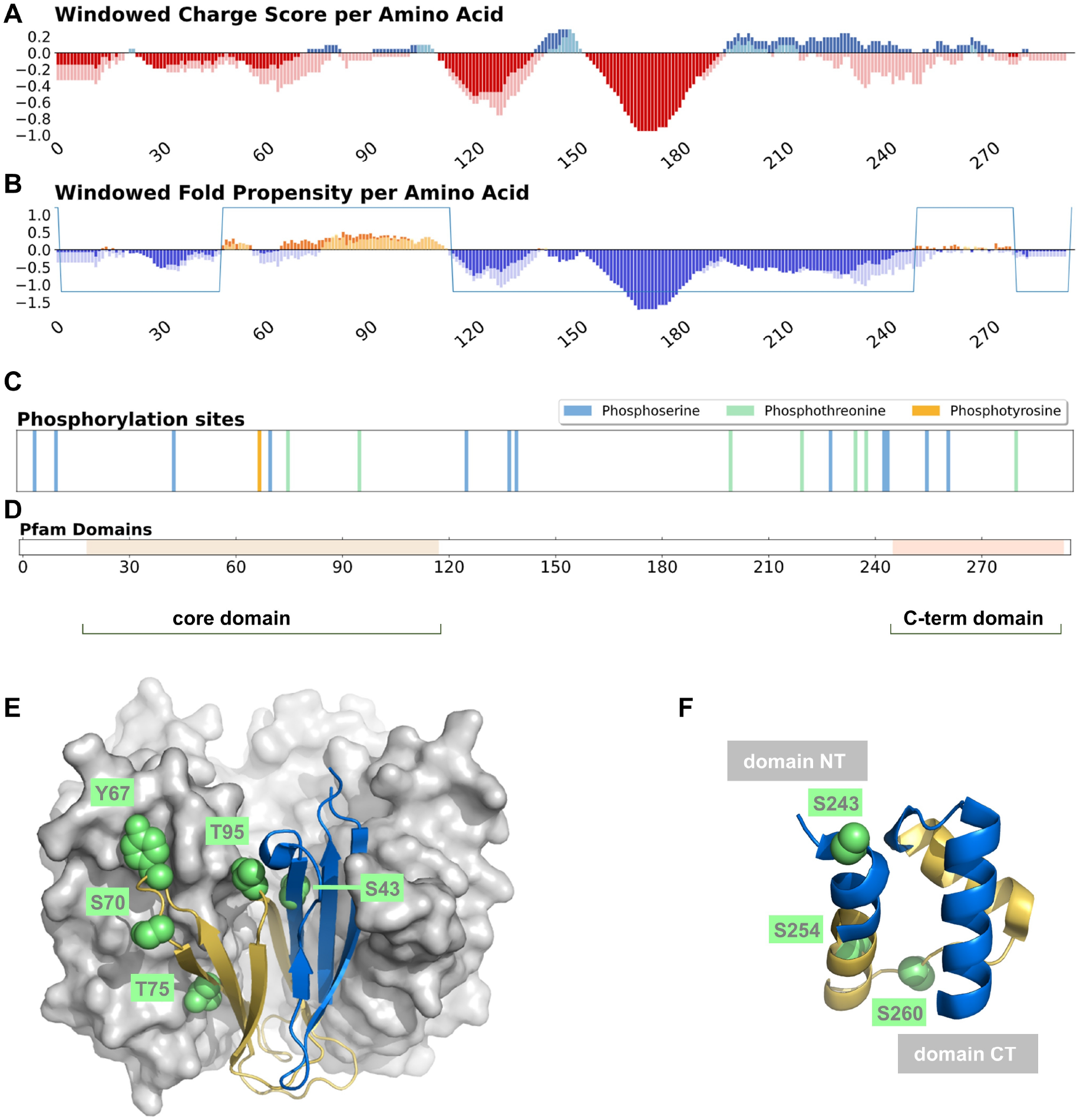
Sequence and structure of human nucleophosmin (UniprotKB: P06748). Windowed charge scores (A), predicted fold propensities (B), phosphorylation sites (C), and Pfam domains (D) are shown, for unmodified and phosphorylated nucleophosmin. The Pfam domains correspond with predicted structured domains, for which solved structure is available, and labelled core domain and C-terminal (C-term) domain. (E) Using the same colour-coding as fold propensity prediction (yellow/folded and blue/unfolded), a cartoon of a core domain monomer is shown against a surface of the remaining 4 monomers in a pentamer unit (PDB ^32^ id 4nm8 ^19^). Known phosphorylation sites are indicated with green spacefill and residue numbers. (F) Also with the yellow and blue colour-coding for predicted structural order from the phosIDP server, the C-terminal domain is drawn (first model from 2vxd ^21^). Known phosphorylation sites for this sequence are shown, along with an indication of the N- and C-terminal ends of the domain.

The other domain known to be structured for nucleophosmin, and also detected with the phosIDP sequence analysis, lies at the C-terminus (CTD, Fig. 2 panels B, D, F). The CTD is predicted as relatively weakly folded, with colour-coding transferred from sequence prediction to the NMR structure (2vxd ^21^), consistent with observation that its folding stability is significantly lower than that of the pentamer unit formed by the core domain, as measured by thermal and chemical denaturation ^22^. Indeed, mutation of the CTD has been associated with loss of structure and amyloid formation, possibly related to pathogenesis ^23,24^. These results again indicate the utility of the server for identifying regions that are predicted to be of relatively low folded state stability. With regard to phosphorylation, notable amongst CTD sites (Fig. 2F) is the location of Ser243 at the start of the first helix in the CTD, and lying in the portion predicted (from sequence, coloured blue) to have only marginal folded state stability. Since favourable interactions arising from a phosphorylated helix N-cap are greater than any amino acid N-cap ^25^, it is possible that phosphorylation at Ser243 is coupled with CTD folding stability and thereby with biological activity.

### Charge distributions and phosphorylation in stress granule proteins

Whilst proteins termed core, such as CIRBP and nucleophosmin, are central to SG formation, client proteins can be found in SGs but do not themselves cause formation of SGs ^14^. Having found in the two examples studied that SG core protein phosphorylation is extensive, we wondered how core protein charge compares with that of client proteins. In order to study protein regions rather than the overall charge of the protein, and examine local effects, charge in overlapping 30 amino acid windows was summed. Further this was divided into regions predicted to be structured or intrinsically disordered, according to the fold propensity results of the server. Distributions of the windowed net charge values are shown for IDRs of core and client proteins, and their phosphorylated counterparts are compared (Fig. 3). Interestingly, unphosphorylated core proteins tend towards overall charge neutrality more than client proteins, but with a distinct enrichment for positive charge. Upon phosphorylation the enrichment for positive charge is largely removed, perhaps indicative (on average) of reduced RNA interaction and a tendency towards SG dissolution more so than formation, although both behaviors have been observed experimentally, dependent on the system ^13^. Although the calculations presented in Fig. 3 are offline, they demonstrate the relevance of studying charge distributions and how they change with phosphorylation.

**Figure 3.**
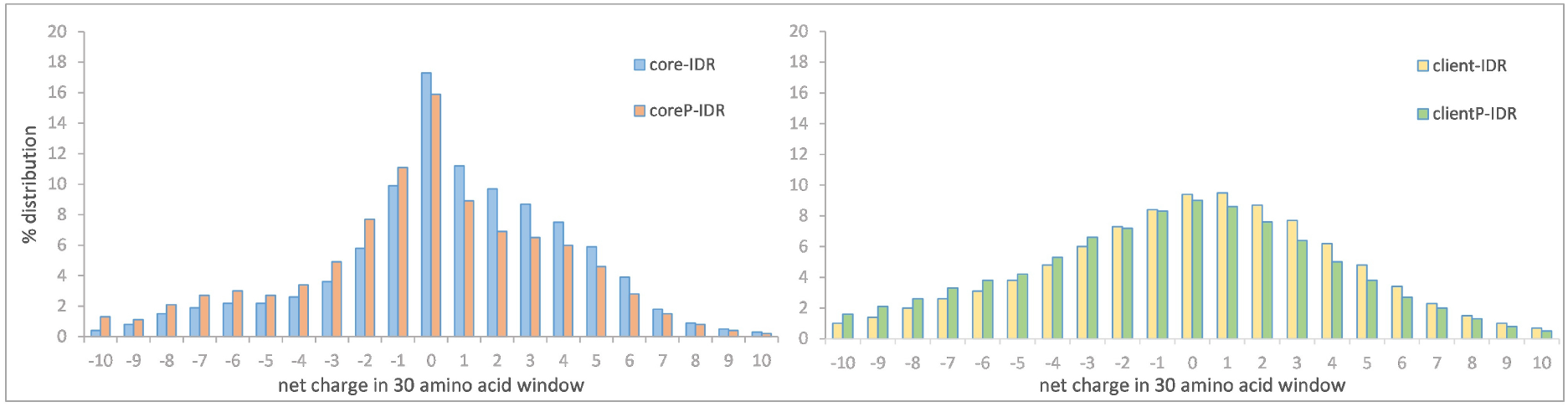
Core and client (and phosphorylated counterparts, denoted P) protein charge distributions, calculated from overlapping 30 amino acid windows within predicted IDRs.

## Discussion

Previous work has demonstrated the importance of net charge and also the balance between positive and negative charge in the analysis of IDR conformation and function ^26^. Here, the crucial role that phosphorylation plays in many systems, not only through site-specific interactions, but also with global switching of charge profiles, can be quickly and conveniently studied with phosIDP, subject to data collation in the UniProt database. Two examples were taken from a dataset of ‘core’ proteins that are integral for membraneless organelle formation. For CIRBP (Fig. 1), phosphorylation of a disordered RGG domain, summed over several sites, reduces the net positive charge close to neutrality, similar to the effect seen overall in IDR regions of the core protein set (Fig. 3). Given the mRNA localisation in membraneless organelles, it is apparent that charge modulation could fine tune stability. In the second example, nucleophosmin (Fig. 2), phosphorylation is enriched close to the boundaries of predicted structured and disordered regions, which for the oligomerisation domain relates to structural transitions that are known to be modulated by phosphorylation. We suggest that the phosIDP web tool will be particularly useful when looking for these regions of predicted weak disorder or structure, and the effect of phosphorylation, which can in turn be matched to available experimental data or lead to experiment design.

## Methods

### Using the PhosIDP web tool

As part of the protein-sol software suite ^15^, PhosIDP is freely available at https://protein-sol.manchester.ac.uk/phosidp without requiring a license or registration. In order for the software to execute the user needs to provide a valid UniProt ID ^16^. Results are displayed graphically with additional information provided below the plots. Data is also available to download as text. The custom URL is available for 7 days from the day of creation.

### The phosIDP algorithm

Upon receiving a UniProt protein ID ^16^, phosIDP calculates charge at neutral pH as a sequence-based profile, averaged over a sliding window of 21 amino acids ^15^, with a second charge profile added that corresponds to the addition of a double negative charge for each phosphorylation site recorded in UniProt (Fig. 1A). A hydropathy scale ^27^ is combined with net charge to predict IDRs ^4,28^, in a scheme repurposed from the existing implementation on protein-sol ^15^. The scale for fold propensity (Fig. 1B) extends from positive/folded to negative/unfolded. An additional feature of the folded state prediction is a sawtooth window that smooths structured and intrinsically disordered regions. Short stretches of IDR within ordered regions may legitimately describe the behavior of loops, but could also obscure the overall domain structure. The sawtooth envelope is created from the predicted profile with the caveat that disordered regions with fewer than 10 amino acids in length take on the average prediction for a window of up to 41 amino acids centered on that region. Additionally, predicted IDRs separated by structured regions of fewer than 10 amino acids are combined into one longer IDR. For reference, phosphorylation sites, as curated by UniProt (Fig. 1C), and Pfam domains ^29^ (Fig. 1D) are displayed.

### Datasets of proteins in membraneless organelles

Three publically available databases were studied in order to generate a dataset of proteins that have been located in membraneless organelles. MSGP ^30^ is a database of proteins in mammalian stress granules, and PhaSepDB ^31^ contains proteins that undergo LLPS in various organelles. Each of these have overlap with DrLLPS ^14^, with the advantage of DrLLPS recording proteins as core, client, or regulator. Since our aim was to compare core and client proteins, DrLLPS was used to generate sets of 56 core proteins and 728 client proteins.

## Data availability

The reported web tool is freely available online. The data associated with LLPS proteins can be provided upon request.

## Acknowledgements

We thank members of the Warwicker and Verma groups for providing feedback on the web page content and layout. The authors would like to acknowledge that this work has been supported by the University of Manchester and the Agency for Science, Technology, and Research (A*STAR) Singapore, and by the UK EPSRC (grant EP/N024796/1). CSV thanks A*STAR for grants (grant IDs H17/01/a0/010, IAF111213C, H18/01/a0/015).

## Author contributions statement

STN, MH, OJ, and JW conducted the studies and analysed the results. All authors were involved in conceiving the work and reviewing the manuscript.

